# Dual autogenous control of the multiple antibiotic resistance phenotype in *Escherichia coli*

**DOI:** 10.1101/008169

**Authors:** Guillermo Rodrigo, Djordje Bajić, Ignacio Elola, Juan F. Poyatos

**Affiliations:** Institute of Systems and Synthetic Biology, CNRS-UEVE, 91030 Évry, France; Centro Nacional de Biotecnología, CSIC, 28049 Madrid, Spain

## Abstract

Bacteria can defend against diverse antibiotics by mounting a multiple antibiotic resistance (*mar*) phenotype. The resistance is linked to a chromosomal locus that encodes an activator and a repressor regulating their own expression. Here, we investigated how this dual autogenous control determines the dynamics of the response. We found that the regulatory architecture provides a mechanism to enable rapid induction, generate pulses of activation, and increase the range of sensing. The response is also graded and homogeneous across the population. Moreover, the interaction of a third regulator with the core module fine tunes the previous features, while limiting the cross-talk with metabolic signals. A minimal model accurately anticipates these properties, and emphasizes how specific attributes of the circuit components constrain the appearance of other potential behaviors associated to the regulatory design. Our results integrate both molecular and circuit-level characteristics to fully elucidate the dynamic emergence of the *mar* phenotype.

## Introduction

The plastic expression of alternative phenotypes enables *Escherichia coli* to adjust to many environmental circumstances (Smits et al., 2006). When these circumstances get particularly adverse for survival, the changes in phenotype normally involve a number of physiological responses that help the bacterium to defend against the effects of the stress (Storz and Hengge, 2011). Timing, strength, and variability of the response become thus essential properties to regulate.

One of these physiological programs corresponds to the <u>m</u>ultiple <u>a</u>ntibiotic <u>r</u>esistance (*mar*) phenotype, which capacitates bacteria to tolerate several toxins including antibiotics like tetracycline or chloramphenicol (George and Levy, 1983). That this response connected for the first time antibiotic resistance to the bacterial chromosome, rather than being caused by a plasmid-borne gene, prompted the search for a better understanding of its genetic architecture. In this way, we currently recognize that the *mar* phenotype is coupled to a unique operon architecture harboring a repressor (MarR) and an activator (MarA), and that it is additionally modulated by other transcriptional factors (e.g., SoxS or Rob) (Cohen et al., 1993a; Cohen et al., 1993b; Martin and Rosner, 2002). Early experiments also identified salicylates (and other repellents) as potential inducers of the phenotype (Rosner, 1985; Cohen et al., 1993b).

The expression of the operon is then consequence of the inactivation of the repressor –MarR represents the sensor of the stress– and the later boost in activator levels –MarA works as the actuator of the system. Increase of MarA abundance act subsequently on a relatively large regulon that includes genes contributing to efflux pumps, e.g., *acrAB-tolC* (Ma et al., 1995; Fralick, 1996), membrane permeability systems, e.g., *micF-ompF* (Cohen et al., 1989), etc. Beyond all molecular details, very little is known about the *dynamic* aspects of the *mar* response, and how these aspects are ultimately determined by the particular genetic circuit that governs its action.

This circuit incorporates several feedbacks (Figure 1A) involving a crucial combination of both negative and positive autogenous control (Goldberger, 1974; Savageau, 1974). Notably, autogenous control was shown to provide very suitable features for stress response (Camas et al., 2006; Smits et al., 2006), such as speedup of dynamics (Savageau, 1974; Rosenfeld et al., 2002) or diversification of cellular behavior (Maeda and Sano, 2006; Ji et al., 2013). How both types of control act together and the consequences of this combined regulation for the mounting of the antibiotic resistance remains, however, an open question.

**Figure 1.**
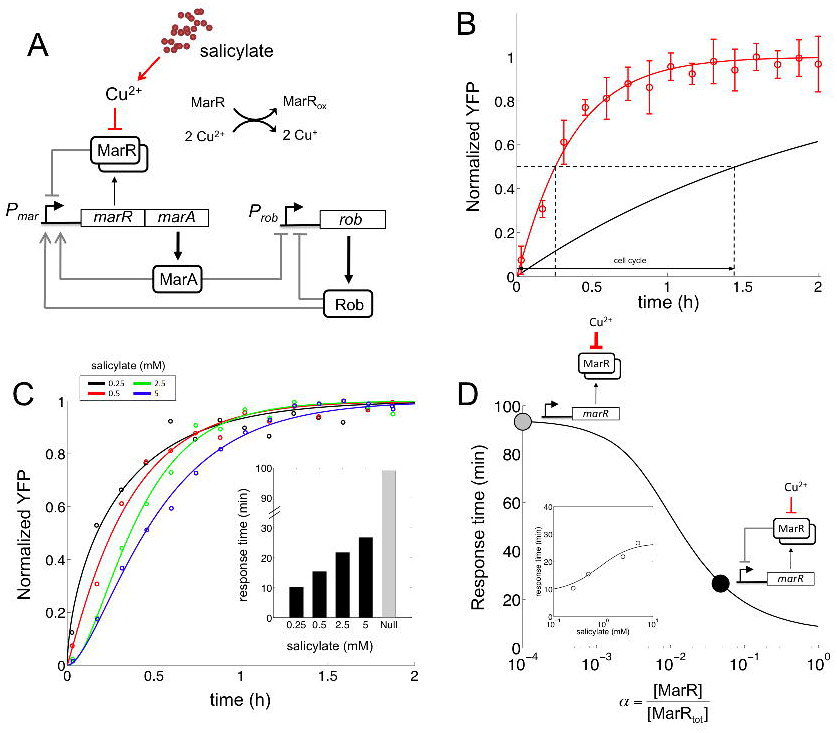
Regulatory architecture and fast dynamic response of the *mar* circuit. (A) MarR dimers, MarA, and Rob regulate the *mar* promoter (P*_mar_*). The response is induced when salicylate causes the intracellular accumulation of Cu^2+^ to oxidize MarR, what prevents its promoter binding. (B) Dynamic response upon induction with 0.5 mM salicylate (error bars correspond to the mean and standard deviations of three independent replicates; fluorescence values normalized by the maximum). The red line corresponds to the fit to (1 − *e*^−^*^λt^*)*^m^* with *λ* = 2.92 ± 0.31 h^−1^ and *m* = 1.08 ± 0.15. This gives a response time (time to reach half of the steady state concentration) of *t*_50_ = 15.35 ± 3.87 min. This time is faster than the cell-cycle time (*t*_cell-cycle_ = 86.64 ± 5.42 min, obtained from the experimental growth curves, *µ* = 0.48 ± 0.03 h^−1^), which represents *t*_50_ of a null model that would assume constant transcription rate. (C) Normalized response of the system to four different concentrations of salicylate. Experimental values (small circles represent averages of three replicates) and theoretical models [lines are the fit to (1 − *e*^−^*^λt^*)*^m^*] are shown. The inset shows the corresponding *t*_50_ values (gray bar denotes response time of the null dynamics, *t*_cell-cycle_ = 99.02 ± 9.43 min, which was calculated with the growth rate at 5 mM salicylate, *µ* = 0.42 ± 0.04 h^−1^). (D) Model predictions of how *t*_50_ would change as a function of the remaining fraction of non-oxidized MarR (*α*) in saturating salicylate conditions (10 mM). The black point corresponds to the predicted value with *α* ≃ 5%, which we estimated from experimental data (Supplemental Information), while the gray point describes a limiting regime in which most MarR molecules are titrated (i.e., no repression). The inset shows, with that value of *α*, a good correspondence between model predictions and experimental values of the response to different dosages of salicylate.

To investigate this issue, we examined the functioning of the circuitry associated to the *mar* phenotype by combining quantitative measurements in *E. coli* with mathematical modeling. We thus distinguished those factors influencing both response time and range. The analysis underlined the role of copper signaling in the dynamics, and explored the consequences of alternative genetic implementations of dual autogenous control. We additionally studied how different (stress) signals are integrated in the *mar* promoter, and the degree of heterogeneity in the response by means of time-lapse microscopy. All these dynamical properties become instrumental to unveil several –yet unknown– functional roles of certain molecular determinants of the phenotype (e.g., Rob), and to better comprehend the circuit-level design principles that eventually orchestrates the response.

## Results

### MarR gives rise to a fast response of the *mar* circuit

To examine the time scale of the activation of the *mar* phenotype, we used a chromosomally integrated yellow fluorescent protein (YFP) under the control of the *mar* promoter (P*_mar_*-*yfp*). We monitored YFP dynamics at high resolution in reaction to salicylate, a phenolic compound known to trigger the response of the system (Figure 1B), and quantified the time to reach half its steady state as response time (*t*_50_). With 0.5 mM salicylate, induction of the phenotype is carried out in about 15 min. This is almost six times faster than the cell-cycle time (which we measured experimentally, Figure S1A). Note that this reference time corresponds to a null model in which we can conceive the phenotype to be under the control of a simple regulation (i.e., non-autogenous control). That leads to present a regular exponential increase [(1 − *e*^−^*^µt^*), with log(2)*/µ* denoting the cell-cycle time] to reach the induced steady state.

The negative autoregulation of MarR could be the primary responsible element for the speedup (Savageau, 1974; Rosenfeld et al., 2002; Camas et al., 2006). This being the case, we expected that *t*_50_ would change with the amount of salicylate, as dosage eventually determines the number of free MarR molecules (i.e., the extent of the feedback repression, Figure S1B); note that MarR is a strong repressor (Seoane and Levy, 1995). Indeed, the higher the dosage the slower the observed response (Figure 1C). Interestingly, we observed that the behavior of the system was still far (in terms of response time) from that predicted without negative autoregulation at saturating levels of inducer, in which free MarR molecules should be mostly absent. We hypothesized that this could be linked to the particular action of salicylate, which inactivates MarR by shifting the intracellular redox equilibrium of copper from Cu^2+^ to Cu^+^ (the system is not induced in anaerobic conditions when this balance is potentially modified, Figure S1C). More in detail, Cu^2+^ oxidizes a residue of MarR that causes tetramerization and repressor dissociation from the *mar* promoter (Figure 1A) (Hao et al., 2014). If this mechanism could not ultimately titrate all repressor molecules, it would impose a mandatory reduction of response time.

We included this effect explicitly in a mathematical model of the circuit that also incorporates the specific regulatory architecture (Figure 1A; see Supplemental Information for detailed description of the model, and Figure S1D for an associated parameter sensitivity analysis; nominal model parameters shown in Table S1). Simulation of the dynamics confirms how the fraction of non-oxidized MarR at saturating concentrations of inducer (*α*) eventually modulates response time. This is illustrated in Figure 1D (and also in Figures S1E-F), where we represented how response time is affected by this parameter (accounting for different intracellular concentrations of Cu^2+^). By using recent data on the action of Cu^2+^ on MarR (Hao et al., 2014), we estimated that the specific value of *α* should be around 5% (Supplemental Information and Figure S1G). This value predicts well the *t*_50_ times observed experimentally (inset in Figure 1D).

### Rob, MarA and MarB further modulate the response

Other molecular components related to the *mar* circuit could affect its response time (Figure S2A). In particular, we first investigated the influence of Rob, the main transcriptional cross-talk experienced by the system under physiological conditions (Chubiz et al., 2012). To this aim, we constructed a Δ*rob* strain in the previous P*_mar_*-*yfp* background. The response of the Δ*rob* strain is still fast compared to the cell-cycle time, but slower than the wild-type system [*t*_50_ = 33.64 ± 4.25 min (Δ*rob*) *vs. t*_50_ = 26.65 ± 3.99 min (wild-type), *U*-test *p <* 0.01, Figure 2A]. This difference in dynamics is linked to the effect of Rob on the transcription rate of the *mar* promoter. Since the presence of Rob can be understood as an enhancing effect on promoter activity (Supplemental Information and Figures S2B-D), its absence necessarily causes a slower response (simulations confirm how response time increases by weakening the strength of the promoter).

**Figure 2.**
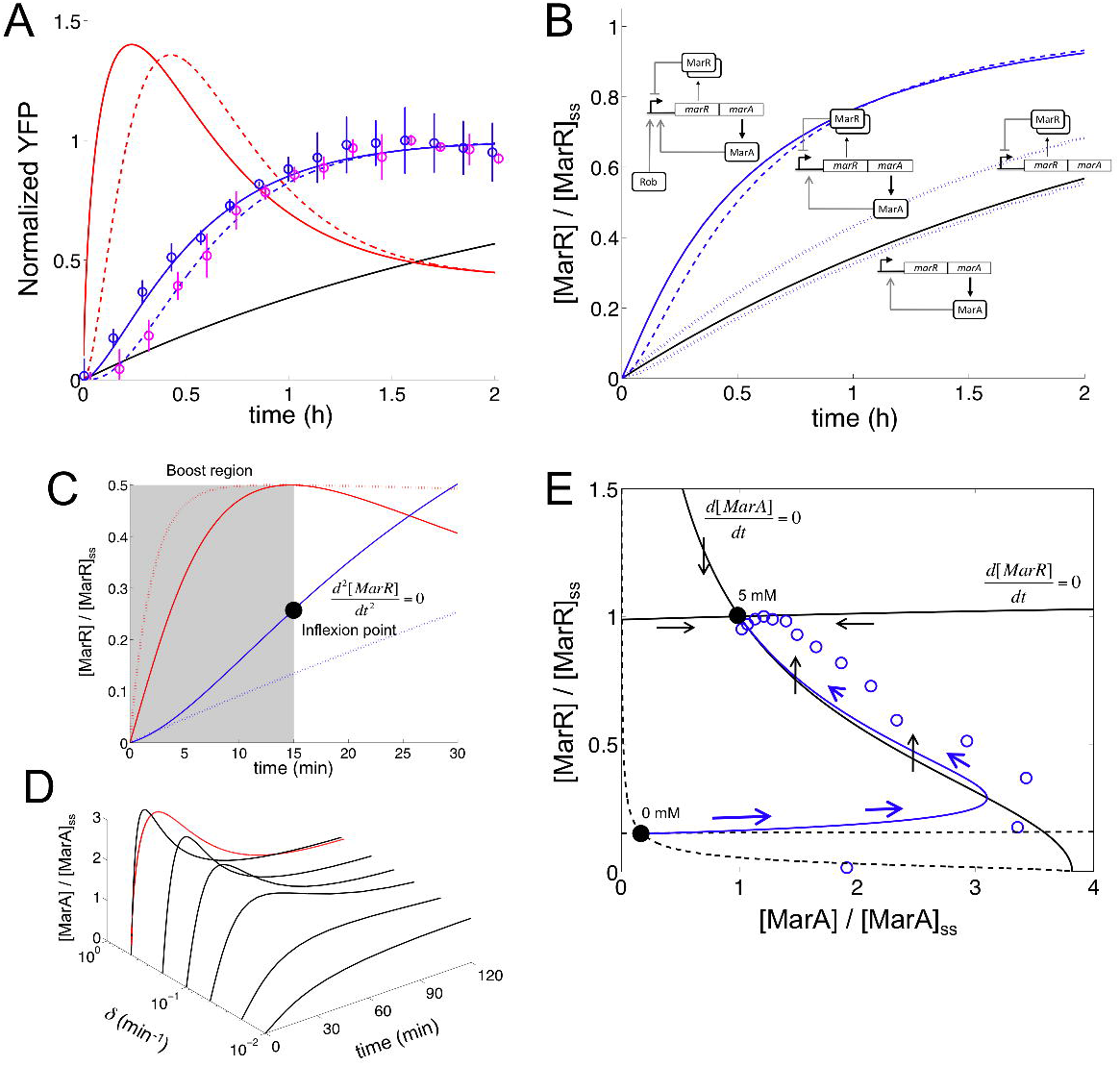
Molecular determinants of the *mar* circuit response. (A) Normalized response upon induction with 5 mM salicylate for the Δ*rob* system (purple circles; mean and standard deviations of three independent replicates; fluorescence values normalized by the maximum). We fitted this data to (1 − *e*^−λ*t*^)*^m^* with *λ* = 2.81 ± 0.68 h^−1^ and *m* = 2.99 ± 1.39 (dashed blue line). We then obtained the corresponding promoter activity (dashed red line). This was compared to both the wild-type strain (*λ* = 2.42 ± 0.38 h^−1^ and *m* = 1.66 ± 0.41; blue circles and solid blue line) and the simple transcriptional unit (black line, *µ* = 0.43 ± 0.01 h^−1^). Δ*rob* response is slightly slower than the wild-type. (B) Model simulations of MarR dynamics [normalized by the steady state (ss) value] upon induction with 5 mM salicylate for alternative regulatory implementations to control the *mar* phenotype. We compared dual autogenous control with (wild-type, solid blue line) or without Rob activation (Δ*rob*, dashed blue line), single autogenous control (positive or negative, dotted blue lines), and no regulation (black line). All simulations performed by using nominal parameter values (Supplemental Information and Table S1). (C) Controlled comparison between the dynamics of circuits with dual (solid line) or negative autogenous regulation (dotted line) at short times (Δ*rob* genotype). Note the inflexion point (black point) where the dynamics changes its curvature and promoter activity is maximal (red line, normalized to help visualization; dotted red line corresponds to the negative autoregulation), while the highlighted (boost) region associates to increasing promoter activity. (D) Model simulations of the MarA concentration (relative to steady state value) upon induction with 5 mM salicylate for different degradation rates of this protein (*δ*). When the protein is unstable, it presents a pulse-like dynamics as it can *follow* promoter activity (red line denotes experimental promoter activity calculated for the wild-type system). (E) Two-dimensional phase space associated to ([MarA], [MarR]) dynamics. Nullclines (black curves; solid for 5 mM and dashed for 0 mM) represent the trajectories in this space where only the concentration of MarR (*d*[MarA]*/dt* = 0 nullcline) or MarA (*d*[MarR]*/dt* = 0 nullcline) changes. The steady states (black points) are then given by the intersection of these curves. We represented a trajectory upon induction with 5 mM from the non-induced state (0 mM); solid line is the simulation and circles the experimental data [obtained from (A), normalizing by the steady state values of YFP and promoter activity]. Blue arrows represent direction and strength of change (the bigger the arrow the faster the change). Black arrows represent direction of change in the 5 mM nullclines.

Moreover, the genetic architecture of the *mar* operon establishes two additional regulatory loops. First, it presents a weak positive feedback implemented by MarA (Martin et al., 2008). This leads (together with MarR) to a combination of positive and negative autoregulation that speeds up the response (compared to the negatively autoregulated system –Δ*marA*, Figure 2B). Indeed, MarA contributes to increase on the fly the strength of the promoter. The combination of regulations originates as well pulses in promoter activity (Figure 2A; we calculated promoter activity by combining independent data of YFP and cell growth dynamics). This is a consequence of the difference in time scales between fast activation [MarA is actively degraded by the Lon protease (Griffith et al., 2004)] and slow repression [MarR presents a low translation rate (Martin and Rosner, 2004)] that eventually originates an inflexion point in activity (Figures 2C and S2E). Regardless of these properties, MarA and MarR have similar (protein) expression levels (quick degradation compensates inefficient translation). Note that the pulse-like behavior in promoter activity is reflected in the dynamics of MarA due to its short half-life (Figure 2D; we considered the experimental concentration of MarA proportional to promoter activity). This also accentuates the specific geometry of the corresponding two-dimensional phase space [associated to the (MarA, MarR) variables, Figures 2E and S2F-I]. Second, the *mar* operon contains yet another open reading frame, MarB (Cohen et al., 1993a). MarB was recently identified as a repressor of the *mar* promoter (Vinué et al., 2013; Nichols et al., 2011) what determines a new putative negative feedback into the system. As expected, we observed experimentally that a Δ*marB* strain exhibits a slower response to salicylate (Figures S2J-N). Overall, our results indicate that Rob, MarA and MarB are regulatory elements that enhance the fast mounting of the *mar* phenotype.

### The *mar* circuit presents wide input and moderate output dynamic ranges

We extended the characterization of the *mar* phenotype with the quantification of the input (*R_in_*) and output (*R_out_*) dynamic ranges of the regulatory module (Goldbeter and Koshland Jr, 1981). To this aim, promoter activity was (experimentally) measured in steady state for different concentrations of salicylate. We then fitted the curve to a sigmoidal to obtain *R_in_* = 31.11 ± 2.46 and *R_out_* = 8.85 ± 0.04 for the wild-type system (inset in Figure 3A). This regime of values, which are captured by our model (Figure S3A), demonstrates that dual regulation increases the sensitivity of the response with respect to a circuit without feedback, in a qualitatively similar manner to negative autoregulation (Figure S3B) (Nevozhay et al., 2009; Madar et al., 2011).

**Figure 3.**
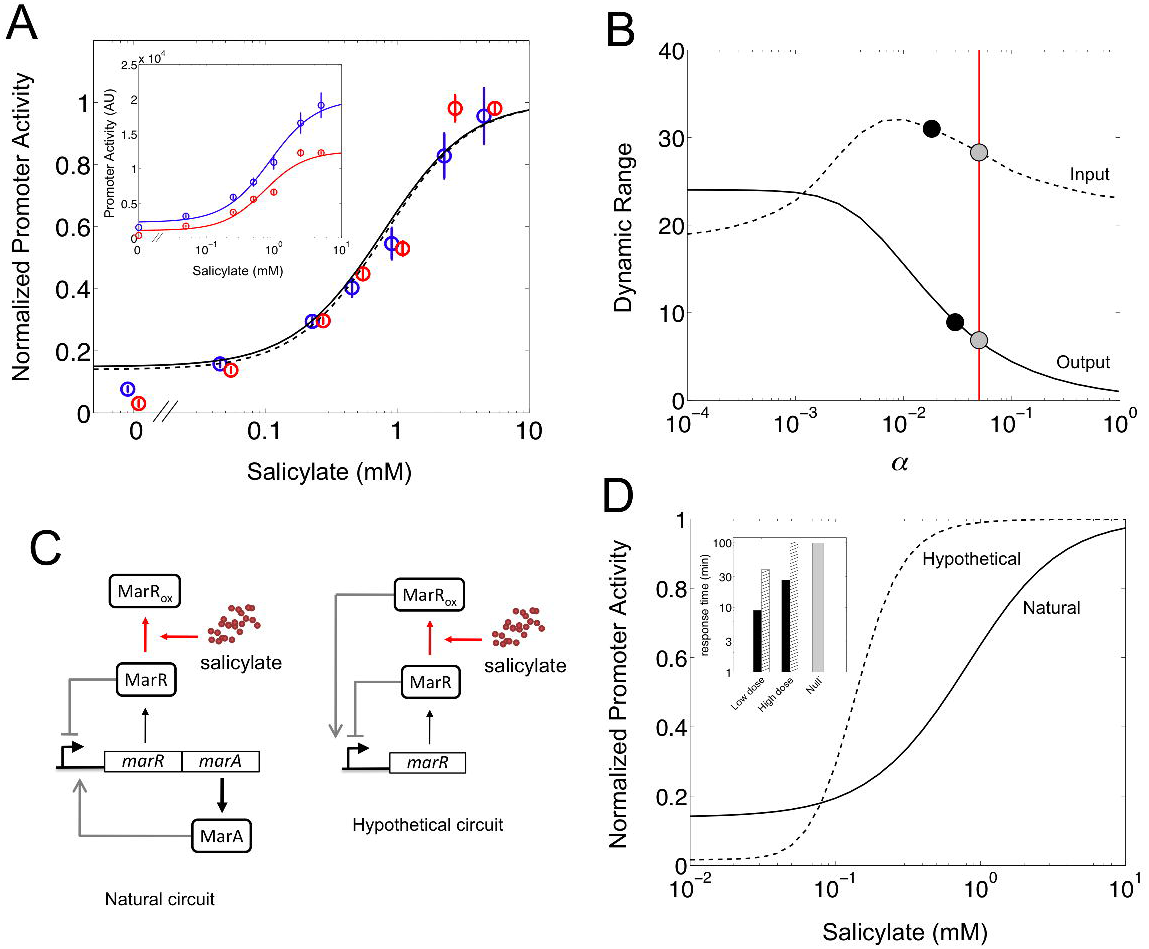
Input/output dynamic range of the *mar* circuit response. (A) Normalized promoter activity in steady state (relative to the maximum) is represented for each concentration of salicylate. Open circles and error bars correspond to experimental data (blue for the wild-type and red for the Δ*rob* systems), while solid and dashed lines correspond to model predictions (wild-type and Δ*rob*, respectively). The inset shows promoter activity in steady state [arbitrary units (AU)]. A Hill-like model was fitted independently to obtain the input and output dynamic ranges [*R_in_* = 31.11 ± 2.46 and *R_out_* = 8.85 ± 0.04 (wild-type), *R_in_* = 20.88 ± 1.17 and *R_out_* = 11.48 ± 1.03 (Δ*rob*); see also Supplemental Information]. (B) Model simulation of the change in input (dashed) and output (solid) dynamic ranges of the system with the remaining fraction of non-oxidized MarR upon induction with salicylate (*α*). Black points correspond to the experimental values reported for the wild-type system, whereas gray points correspond to the model predictions for the chosen parameterization. The value of *α* used in this work (*α* ≃ 5%), inferred from the oxidation curve with copper, is marked by a vertical red line. (C) Circuit schemes of two possible molecular implementations of dual autogenous control (see Table S2). (D) Model predictions of normalized promoter activity with salicylate. The solid line corresponds to the *mar* circuit with Δ*rob* (*R_in_* = 27.51 and *R_out_* = 7.17), whereas the dashed line corresponds to a hypothetical circuit where the oxidized MarR acts as an activator (*R_in_* = 5.16 and *R_out_* = 62.17). The inset shows the corresponding response times of these two implementations of dual autogenous control. Black bars correspond to the natural circuit, whereas hatched bars to the hypothetical circuit (gray bar is for the null model). Low dose corresponds to 0.01 mM and high dose to 10 mM.

Rob appears not to change much the previous input/output ranges but rather to scale the response according to simulations of our model (Figure S3C). The scaling factor can be approximated by *ρ*^1^*^/^*^3^, with *ρ* denoting the activation fold by MarA and Rob (see Supplemental Information). To evaluate these predictions, we produced a experimental dose-response curve of the Δ*rob* strain and fitted it again to a sigmoidal. The data confirmed that absence of Rob does not alter much the range values (*R_in_* = 20.88 = ± 1.17 and *R_out_* = 11.48 ± 1.03), but scaled the expression of the system (the wild-type dose response is about 1.6-fold that of the mutant). Indeed, the fold change observed (1.6^3^ ≃ 4) agrees with experimental estimations of the activation fold by MarA (Martin et al., 1996; Chubiz et al., 2012) (see Figure S3D for other experimentally obtained fold-change values that are captured by our model).

We then asked if the remaining non-oxidized MarR (upon induction with salicylate) influenced dynamic ranges as it did previously with response time. Figure 3B shows the (model) predictions of the change in *R_in_* and *R_out_* with *α*, together with the experimental values reported in this work. Note that the output dynamic range follows a similar dependence with *α* as the response time (i.e., the higher is the derepression of the system, the higher and slower is the induction). A fast response then exhibits moderate *R_out_*. This trade-off is nicely captured by solving analytically the model, giving *R_out_* = *α*^−2^*^/^*^3^ and *t*_50_ ∝ *α*^−1^ (Supplemental Information). We could then calculate in an alternative manner *α* from the experimental value of *R_out_*. We obtained a value of *α* ≃ 4%, which corroborates the value inferred from the oxidation curve with copper (*α* ≃ 5%). In contrast to the monotonous trend of *R_out_* with *α*, *R_in_* presents a maximum although not very pronounced.

### The polycistronic implementation of dual autogenous control shows higher sensitivity and faster response

An alternative genetic implementation of dual autogenous control could involve dual regulators. For instance, in *E. coli* there exist three systems regulated by this type of architecture: CRP, ChbR, and LldR. These transcription factors work as repressors that turn into activators in response to cAMP (Ishizuka et al., 1994), chitobiose (Plumbridge and Pellegrini, 2004), and lactate (Aguilera et al., 2008), respectively. Based on these cases, we imagined an *hypothetical mar* circuit in which the oxidized MarR functions as an activator (Figure 3C). We modified accordingly our model to solve the new dynamics (Supplemental Information). The natural circuit showed higher *R_in_* (approximately 5-fold to the hypothetical one), whereas the hypothetical circuit exhibited much higher *R_out_* (about 9-fold to the natural one, Figure 3D).

This suggests that the specific architecture of the *mar* circuit, which is unique in the *E. coli* genome (Table S2), preferentially evolved to exhibit higher sensitivity to gradients of pollutant concentrations, while maintaining moderate fold change. Alternatively, responses governed by dual regulators could require a wider output range to either fine tune very large regulons (this applies for instance to CRP; note that the *crp* and *mar* modules are connected, Figure S3E) or induce fairly digital responses [e.g., to activate the response under a very narrow signal range, as it is the case of ChbR, which activates the expression of the *chb* operon only when sufficient flux through their associated pathway is sensed (Plumbridge and Pellegrini, 2004)]. In addition, the response dynamics of the natural circuit would be faster than the hypothetical one (inset in Figure 3D, and also Figure S3F). More in detail, we found that the hypothetical system would respond similarly to one under simple regulation at high dosages (activation would dominate repression) but exhibit speedup at very low dosages. In the latter situation, there would be a combined action of positive and negative regulation over the promoter, similar to the one governing the induction of the *mar* circuit.

### Rob reduces the cross-talk between antibiotic and metabolic stresses

The regulation of the *mar* promoter by additional transcription factors makes it responsive to other signals (Ruiz and Levy, 2010) like oxidative stress (through SoxS), DNA supercoiling (through Fis), or metabolic/catabolic stress (through CRP, Figure 4A). Since cAMP-CRP signaling is instrumental to coordinate gene expression in multiple situations (You et al., 2013), we decided to analyze how the promoter integrates both the antibiotic and metabolic signals as a two-dimensional transfer function (Kaplan et al., 2008). Because the *mar* and *crp* operators overlap, we extended our model by considering competitive binding between the activators MarA, Rob and CRP. In Figure 4B, we show the promoter activity obtained experimentally (in steady state) for several combinations of salicylate and cAMP (see Figures S4A,C for the dynamics). The response appeared almost independent of cAMP. We observed however a significant effect of cAMP on the Δ*rob* strain (Figure 4C, see also the dynamics in Figures S4B,D). This effect was predicted by our simulations (Figures S4E-F). We could then suggest that the highly expression of Rob (Skarstad et al., 1993) buffers the activation by cAMP.

**Figure 4.**
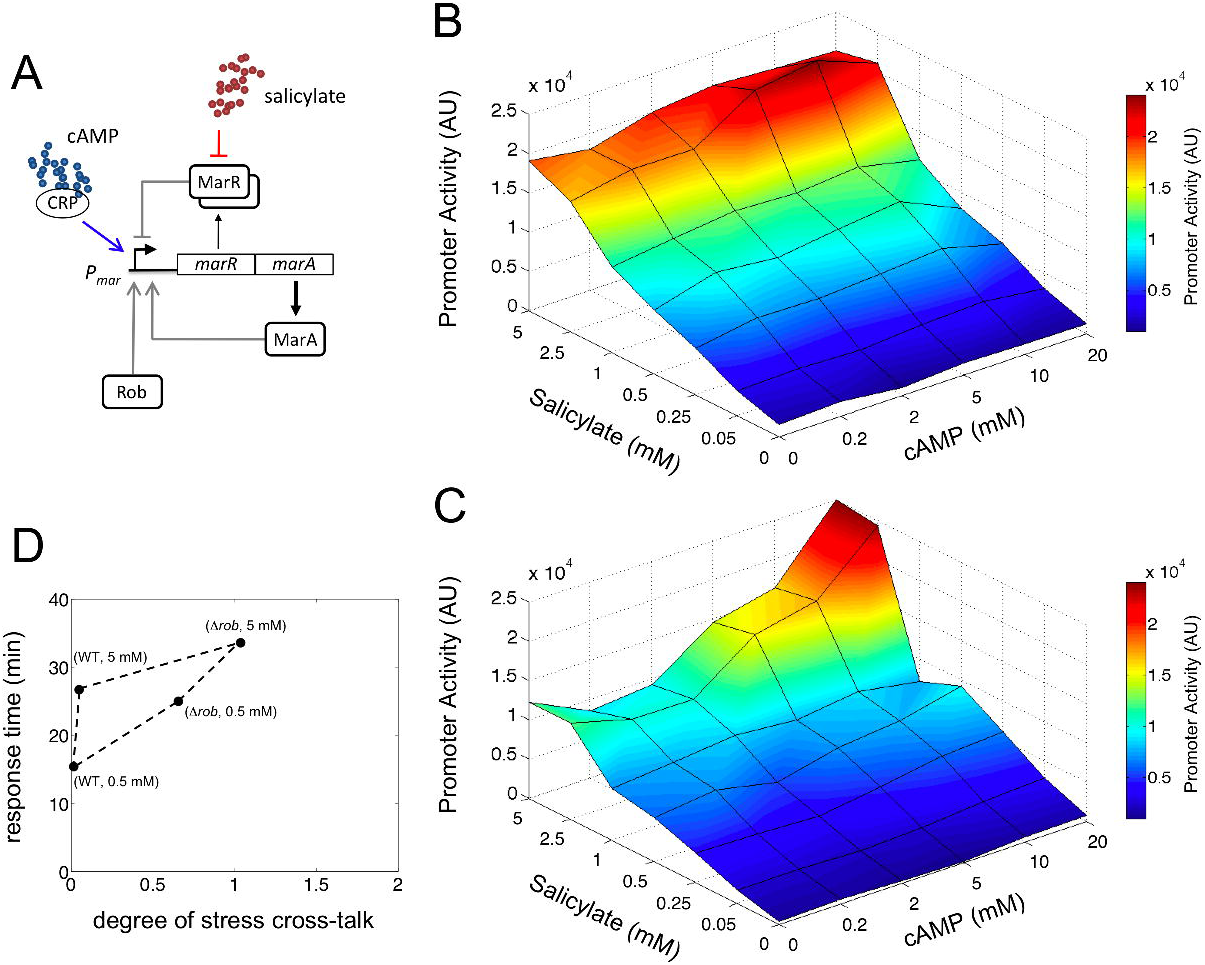
Two-dimensional transfer function of the *mar* circuit response. (A) CRP-cAMP, in addition to salicylate via derepression of MarR, can induce the *mar* promoter. (B, C) Promoter activity in steady state [arbitrary units (AU)] is represented for each of the 42 combinations of the two input signals, salicylate and cAMP. (B) Wild-type system; (C) Δ*rob* system. (D) Representation of the genotype (*rob*) by environment (salicylate) interaction on the phase space defined by the response time and the degree of stress cross-talk (a degree of 1 implies that promoter activity with 20 mM cAMP is 100% higher than without cAMP, see also Supplemental Information).

To link this result with the dynamics of induction, we quantified the cross-talk between both inputs by comparing the promoter activity without or with cAMP (20 mM) for a particular salicylate dosage. Figure 4D shows the genotype (*rob*) by environment (salicylate) interaction, pointing out the double effect of Rob on the system. Thus, Rob appears as a regulatory element that *E. coli* employs, in addition to fine tune response time, to isolate the *mar* phenotype from metabolic signals.

### The *mar* phenotype remains buffered without stress

The regulatory motif associated to the *mar* response could give rise to excitable dynamics –even without any stress signal– due to the presence of positive and negative feedbacks corresponding to different time scales (Hasty et al., 2002; Guantes and Poyatos, 2006). To explore the possible appearance of this behavior, we simulated the stochastic dynamics of the system (Supplemental Information, Figure 5A). The time series presents, at the single cell level, fluctuations out of the (deterministic) equilibrium caused by the mixture of gene expression noise (Munsky et al., 2012) and the action of dual regulation (Figure 5B).

Moreover, that the circuit topology could potentially lead to excitable, or pulsing, dynamics can be further contemplated by modifying some of its basic attributes. Indeed, either a variant model that did not include MarA as monomer (i.e., as a multimer, Figures S5A-C) or a different one that incorporated asymmetric and strong competitive binding (of MarA on MarR) exhibited more complex dynamics (Figures S5D-F) (Hasty et al., 2002; Garcia-Bernardo and Dunlop, 2013). Thus, to confirm the predictions of our original model, we followed experimentally the dynamics of a single cell in absence of salicylate. Figure 5C illustrates a representative trajectory that corroborates the noisy dynamics around steady state without the emergence of significant oscillations (much of the cell-to-cell variability in YFP and CFP is correlated, Figures 5D, S5G-H). The histogram of variability on YFP suggests a constant production of the proteins of the *mar* operon and the absence of any oscillatory/pulsing dynamics (Figure 5E) (Munsky et al., 2012).

**Figure 5.**
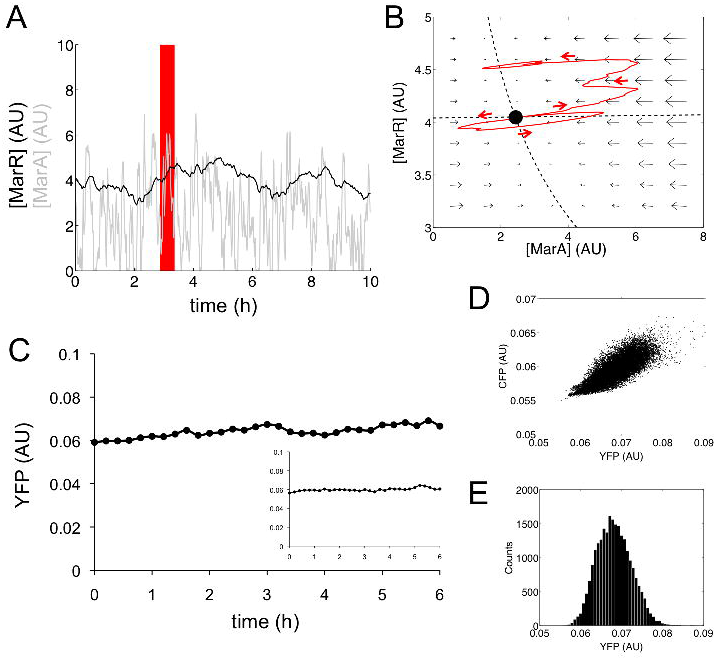
Non-induced stochastic dynamics of the *mar* circuit. (A) Model simulation of the stochastic dynamics of MarR and MarA in absence of salicylate [arbitrary units (AU)]. (B) We plot some part of the trajectory (red box) in (A) in the phase space of the system (red arrows indicate time evolution). This representation highlights how gene expression levels fluctuate around the deterministic steady state (black point) by the combined action of intrinsic/extrinsic noise, and dual autogenous control. Dashed lines correspond to the nullclines of the system (recall Figure 2E), and field arrows denote the strength of change towards the steady state (bigger arrow implies bigger change). (C) Representative trajectory (YFP) of a single cell (wild-type system). In the inset, trajectory for the Δ*rob* system. (D) Scatter plot of single-cell YFP and CFP corresponding to an experiment that follows colony growth and gene expression dynamics without salicylate (wild-type). Note the correlation in the fluctuations in both reporter proteins –expressed from different promoters. (E) Distribution of YFP for all single cells at all time points (wild-type) confirms a unimodal distribution corresponding to a continuous production of MarR.

### The *mar* response is relatively homogeneous across a population

We additionally investigated the single-cell dynamics in the presence of stress by tracking cells upon induction with 5 mM salicylate at time 0 (Figure 6A; CFP signal also shown as control). We asked to what extent the *mar* phenotype is expressed differentially, e.g., only triggered in a subset of the population (Colman-Lerner et al., 2005). By following the dynamics of a growing population of cells, we observed however that activation is relatively homogeneous, with unimodal distributions being clearly identified at different times (Figure 6B, see also Figure S6A), or at various salicylate dosages at steady state (Figures S6B-C). We also noticed that noise decreases with salicylate dosage (Figures S6D-F).

**Figure 6.**
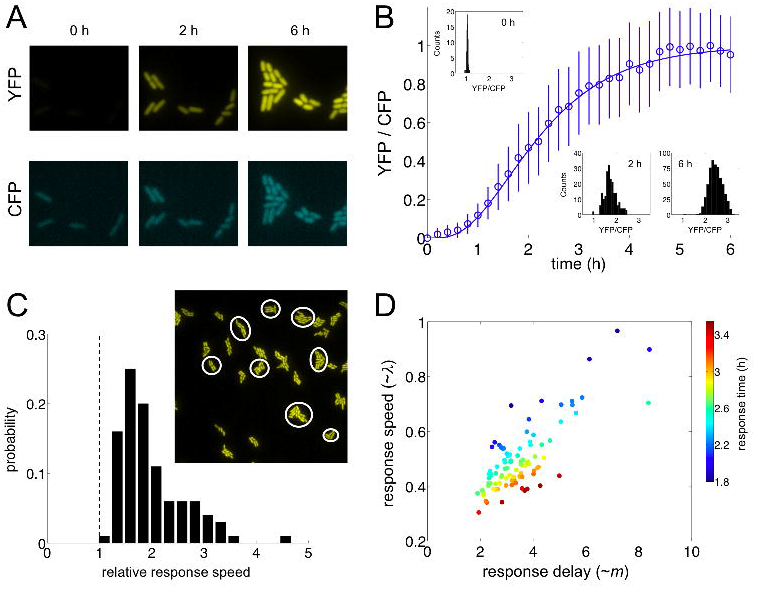
Single-cell dynamic response of the *mar* circuit. (A) Response of the *mar* promoter (YFP signal) and a control promoter (CFP signal) upon induction with 5 mM salicylate at time 0. (B) Graded transcriptional activation of the *mar* phenotype, measured as the ratio YFP/CFP, in a growing population of single cells upon induction with 5 mM salicylate (histograms at different times illustrate the relatively homogeneous response). (C) Variation in relative response speed (as the ratio between response speed and growth rate) among 100 different lineages within the population (see main text). The image shows a subset of lineages. (D) The combination between response speed and response delay ultimately shapes response time. Each point represents an independent lineage.

The reported single-cell behavior further emphasizes that the activation of the phenotype is better understood as a composition of two features: response speed (a measure of how much the expression changes per unit of time) and delay (time before increasing expression, which relates to response nonlinearity, Figure S7A). These properties are determined by *λ* and *m*, respectively, the two parameters characterizing our fit [(1 − *e*^−^*^λt^*)*^m^*, Figure S6G]. By measuring variability in response speed (computed its value at *t*_50_, *µ* for simple regulation and ∼ *λ* for feedback-regulated systems; Supplemental Information), we showed that lineages presented twice the response speed of the simple regulation on average (Figures 6C and S6H-K). Note also that response speed and delay are correlated, and ultimately determine response time (Figures 6D and S6L).

### Discussion

The mechanism of autogenous control of gene expression involves a genetic program in which the protein encoded by the structural gene is working as its own regulatory element. This entails a number of functional advantages with respect to the *classical* model of regulation (Jacob and Monod, 1961; Goldberger, 1974). Notably, these advantages are specific on whether the autoregulation is positive (activator-controlled) or negative (repressor-controlled) (Savageau, 1974). We focused here on an inducible system that presents both types of autogenous regulation (positive and negative) within the architecture that associates to the *mar* phenotype.

Dual control necessarily integrates contrasting properties of the dynamics. For instance, response time could be expected to be either fast or slow according to earlier studies on negative (Rosenfeld et al., 2002; Camas et al., 2006) and positive (Maeda and Sano, 2006) autogenous systems. We observed experimentally a rapid induction that is modulated by the action of salicylate on the strong repression of MarR (Figures 1B-C). Thus, the faster the response, the stronger the repression which also results in lower expression levels at equilibrium (Figure S7B). In addition, the speedup appeared influenced by copper signaling, since the accumulation of intracellular Cu^2+^ in response to salicylate oxidizes MarR preventing its binding to DNA (Hao et al., 2014). Because the intracellular concentration of this cation is bounded (to avoid toxicity), copper balance imposes a maximal repressor abundance for the promoter to be derepressed (Supplemental Information and Figure S1G); a constraint that provides a rationale for the limited translation rate observed for MarR (Martin and Rosner, 2004).

Moreover, the positive autoregulation rather than delaying the response contributes to its speedup due to the increase on the fly of the transcription rate of the promoter (Figure S7C; dynamics of wild-type and Δ*marA* systems shown in Figure 2B). This coordination between repressor and activator generates pulses in promoter activity (Figures 2A and S2B-E), which can be followed by the actuator of the regulon (MarA) due to its short half-life (Griffith et al., 2004). The features that effectively minimized the delay of the response (Figure S7D) are the weak activation and monomeric action of MarA. These attributes maintain the phenotype buffered in absence of signal (Figure 5) (Camas et al., 2006), limit large-amplitude (and heterogeneous) transient responses (Figure 6) (Cağatay et al., 2009), and ultimately set apart the observed dynamics from that characterized in other biological systems also regulated by interlocked positive and negative feedback loops, e.g., (Stricker et al., 2008; Ji et al., 2013).

Interestingly, the essential weakening of the positive control (through MarA) becomes somehow neutralized by the additional regulation of Rob, an interaction that can be effectively interpreted as a mean to increase promoter strength (Figure 3A). The absence of Rob causes not only a slightly slower response, but also a reduction in signal specificity (Figure 4D). While the wild-type circuit decreases the potential cross-talk between antibiotic and metabolic stress signals (the latter connected to the *mar* system through CRP, Figure 4A), the absence of Rob amplifies the transfer between them.

The dual autogenous control of the *mar* phenotype is also unique in its genetic organization. Indeed, this system is the only one in *E. coli* presenting a polycistron carrying both repressor and activator. However, we did identify alternative designs of dual autogenous control by means of dual regulators in this bacterium. These regulators switch between repression or activation in the presence of its cognate inducer [CRP, for instance, works in this way (Ishizuka et al., 1994)]. What are the differences in dynamics between these two implementations? To analyze this, we considered an hypothetical *mar* system in which oxidized MarR turns into an activator (Figure 3C). This more compact genetic design exhibits a smaller input range (i.e., the reaction to a signal gradient is more sigmoidal), a wider output range (what enables, for example, to regulate differentially large regulons as it could be the case of CRP), and a slower response (with no speedup at high dosages) (Figure 3D). This exemplifies that the dynamics corresponding to a given regulatory logic is certainly influenced by its genetic implementation (Guet et al., 2002; Guantes and Poyatos, 2006; Cağatay et al., 2009), and that the *mar* control circuit is quite distintive in its organization and function in order to trigger a response that is fast, graded, sensitive, robust, and coherent across the population.

## Experimental Procedures

### Strains, culture media and reagents

Two-color fluorescent reporter *E. coli* strains (IE01, IE02 and DB01) were engineered to measure the activity of the *mar* promoter. The strain IE01 contains a chromosomal copy of the *yfp* gene under the control of the *mar* promoter, and the *cfp* gene expressed with a constitutive promoter. The strains IE02 and DB01 were constructed by deletion of the *rob* and *marB* genes, respectively, on IE01 with the application of a knockout protocol (Datsenko and Wanner, 2000). Medium LB was always used for overnight cultures. Minimal medium M9 (M9 salts 1x, MgSO_4_ 2 mM, CaCl_2_ 0.1 mM, glucose 0.4%, casamino acids 0.05%, vitamine B1 0.05%) was used to grow cells during characterization experiments. To induce the *mar* circuit, different concentrations of salicylate and cAMP (Sigma Aldrich) were used. When appropriate, kanamycin was used at 50 µg/mL. See also the Extended Experimental Procedures for more details.

### Quantification of fluorescence in a cell population

Cultures (2 mL) inoculated from single colonies (three replicates) were grown overnight in LB medium supplemented with glucose 0.4% at 37 *°*C and 170 rpm. Cultures were then diluted 1:200 in M9 minimal medium and were grown for 2 h at the same conditions. Cultures were then used to load the wells (200 µL) of the microplate (Thermo Scientific) with salicylate and cAMP when appropriate. The microplate was assayed in a Victor X2 (Perkin Elmer) measuring absorbance (600 nm), YFP (497/16 nm, 535/40 nm), and CFP (434/17 nm, 479/40 nm) for 4 h at 37 *°*C with shaking. Analysis of fluorescence data described in Supplemental Information.

### Quantification of fluorescence in single cells

Culture (2 mL) inoculated from a single colony was grown overnight in LB medium supplemented with glucose 0.4% at 37 *°*C and 170 rpm. Culture was then diluted 1:200 in M9 minimal medium and was grown for 4 h at the same conditions. Culture diluted 1:10 (2 µL) was then used to load the agarose pad. Before characterization, 5 mM salicylate was added to induce cells. Agarose pads were monitored in an inverted microscope Axiovert200 (Zeiss) with objective 100X/1.45 oil Plan-Fluar at 30 *°*C. Cell images were acquired from the bright-field and fluorescence channels, YFP (490-510 nm, 510-560 nm) and CFP (426-446 nm, 460-500 nm), using software MetaMorph (Universal Imaging). Analysis of single cell images described in Supplemental Information.

### Empirical modeling the dynamic behavior

A generalized exponential model was used to describe the dynamics of the system upon induction with salicylate, at both population and single cell levels. A sigmoidal model was used to describe the dose-response curve. See more details in Supplemental Information. Parameters for the empirical models were obtained through nonlinear regression with our own experimental data. Bootstrapping was applied to calculate the errors associated to the measurements of response time and input/output dynamic range. *U*-tests were performed to compare distributions of inferred parameters.

### Modeling the *mar* circuit

A system of differential equations was constructed to model the dynamic response of the system. The model considered as variables the concentrations of MarA and MarR. The concentration of Rob was considered constant. Model parameters were mainly obtained from previous experimental data. The model was numerically and analytically solved. The model was perturbed to account for the dynamics of other regulatory architectures. A Langevin approach was followed to simulate stochastic dynamics. See also the Extended Experimental Procedures for more details.

## Supplemental Information

Supplemental Information includes Extended Experimental Procedures, seven Figures, and two Tables, and can be found online.

## Acknowledgements

This work was supported by Spanish Ministry of Economy and Competitiveness grant BFU2011-24691 (JFP), and AXA Research Fund (GR). We thank T. Miyashiro and M. Goulian for strains, J. M. Gómez-Gómez, J. Rodríguez-Beltrán, E. Majer and A. Willemsen for technical assistance, and J. L. Rosner and R. G. Martin for discussions on earlier studies on the *mar* system.

